## Supporting Information for "How *µ*-Opioid Receptor Recognizes Fentanyl"

#### 1 Methods and Protocols

All fixed-charge simulations (weighted ensemble and equilibrium molecular dynamics) were carried out using the GPU-accelerated *pmemd* engine in AMBER18.[S1] The continuous constant pH molecular dynamics (CpHMD) simulations were carried out with the CHARMM program (version c42). [S2] The protein was represented by CHARMM36m [S3] and CHARMM22/CMAP force fields [S4,S5] in the fixed-charge and CpHMD simulations, respectively. Water was represented by the CHARMM-style TIP3P force field.[S2] The POPC and cholesterol molecules were represented by the CHARMM36 lipid force field.[S6,S7] The force field parameters for fentanyl were obtained using the ParamChem CGENFF server.[S8]

**Equilibration simulation of apo active mOR in a lipid bilayer.** The X-ray crystal structure of mOR in complex with BU72 (PDB ID: 5C1M)[S9] was used as the starting model for apo active mOR. The crystal structure represents the wild type but contains a cysteine-s-acetamide (YCM) at position 57. This residue was converted to a cysteine (Cys57). A cholesterol molecule was resolved in the X-ray structure and is bound to the extracellular leaflet near TM7. This cholesterol and all crystal waters in the interior of mOR were kept. Seven additional water molecules were added using the DOWSER program.[S10] Apo mOR was oriented with respect to membrane using the OPM (Orientations of Proteins in Membranes) database.[S11] The CHARMM-GUI web server[S12] was then used to construct the system of mOR embedded in a POPC (1-palmitoyl-2-oleoyl-glycero-3-phosphocholine) lipid bilayer. The disulfide bond observed in the crystal structure was imposed between Cys140 and Cys217. All titratable residues were fixed in the standard protonation states. All histidine residues were set to the neutral $N_\delta$ tautomer (HID) as in the default setting of CHARMM.[S2] The system was first energy minimized using the steepest descent followed by conjugate gradient algorithm. The system was then gradually heated from 0 to 310 K over 200 ps with harmonic restraints on the protein heavy atoms, lipids, and bound water molecules (same as Step 1 in Table S1). Following heating, the system was equilibrated for 117 ns, during which time the various harmonic restrains were gradually reduced to zero (Table S1). The final dimension of the system was $\sim$87$\times$87$\times$113 Å$^3$. The final snapshot was used for constructing the fentanyl-bound mOR model and as the starting structure for the replica-exchange CpHMD simulations of apo mOR (CpH-apo).

**Relaxation of the docked fentanyl-mOR complex in a lipid bilayer.** The fentanyl-bound mOR model was prepared by superimposing a top fentanyl binding pose from a previous docking study[S13] onto the final snapshot from the aforementioned equilibration simulation of apo active mOR. The docked pose showed a salt bridge between the piperidine nitrogen and Asp147 and an aromatic stacking between the phenethyl ring and

His297. The docked model was equilibrated for 115 ns, using harmonic restraints imposed on the protein and fentanyl heavy atoms in the first 5 ns (details see Table S2). During the unrestrained part of the simulation, the root-mean-square deviation (RMSD) of the fentanyl heavy atom positions with respect to the docked structure steadily increases in the first 20 ns and stabilizes at about 6 Å in the remainder of the 110 ns simulation (Fig. S1A). Concomitant with the fentanyl RMSD increase, the minimum distance between fentanyl's piperidine nitrogen and His297's imidazole nitrogen decreases from 10 to about 7 Å (Fig. S1D); however, the salt bridge between the positively charged piperidine amine and the negatively charged Asp147 remains largely stable except for occasional excursions (Fig. S1C), consistent with a previous $\mu$s simulation study.[S14] The final snapshot was used as the starting configuration for the WE simulations.

**Weighted ensemble MD simulations.** Weighted ensemble (WE) is a path sampling protocol that uses splitting and merging trajectories to enhance sampling of rare events. Briefly, the configuration space is divided into bins based on a predetermined progress coordinate and a fixed number of walkers (trajectories) per bin is targeted. At the beginning of the simulation, walkers are initiated from a single bin and after a specified time interval, resampling is performed by evaluating the number of walkers per bin, and for bins with less than desired number of walkers, the walker is replicated (or split), and for bins with more than desired number of walkers, the walkers are pruned. Thus, over time, more bins are sampled and the simulation progresses along the progress coordinate. Details theory and algorithm can be found elsewhere.[S15–S17]

Two WE MD simulations were carried out starting from the equilibrated fentanyl-mOR complex structure, with His297 fixed in either HIE (WE-HIE) or HID (WE-HID) state. The Python-based tool WESTPA[S16] was used to control the WE protocol and data storage. The root-mean-square deviation (RMSD) of the fentanyl heavy atom positions with respect to their starting positions was used as the progress coordinate. The configuration space was divided into bins that covered the RMSD values of 0 and 10 Å. A target number of four and five walkers per bin was used for the WE-HIE and WE-HID simulations, respectively. The fixed time interval for resampling of each walker was 0.5 ns. The bin widths were changed manually in the beginning of the simulations to further accelerate sampling, and the final bin boundaries were placed at the following RMSD values: 0, 0.5, 1, 1.25, 1.5, 1.75, 2, 2.1, 2.2, 2.3, 2.4, 2.5, 2.6, 2.7, 2.8, 2.9, 3, 3.1, 3.2, 3.3, 3.4, 3.5, 3.6, 3.7, 3.8, 3.9, 4, 4.1, 4.2, 4.3, 4.4, 4.5, 4.6, 4.7, 4.8, 4.9, 5, 5.25, 5.5, 5.75, 6, 6.25, 6.5, 6.75, 7, 7.25, 7.5, 7.75, 8, 8.25, 8.5, 8.75, 9, 9.5, 10, $>$ 10. The WE simulations employed the Langevin thermostat, as a stochastic thermostat is required to for the WE strategy to generate continuous pathways with no bias in the dynamics. A total of about 300 iterations were conducted for WE-HIE and WE-HID simulations. From the WE-HID simulation, we uncovered an alternative binding mode that involved a hydrogen bond between fentanyl's piperidine nitrogen and His297's N$\epsilon$. The bound pose was used as the starting configuration for the equilibrium simulation MD-H297(HID). To prepare the starting configurations for the equilibrium simulations MD-H297(HIE) and MD-H297(HIP), the protonation state of His297 in the bound pose was switched followed by 65 ns equilibration (Table S3). The final snapshot which no longer contained the piperidine-H297(N$\epsilon$) hydrogen bond were used to start the MD-H297(HIE) and MD-H297(HIP) simulations.

**Continuous constant pH molecular dynamics (CpHMD)** We applied the membrane-enabled hybrid-solvent continuous constant pH molecular dynamics (CpHMD) method[S18,S19] to determine the protonation states of all titratable residues in the mOR systems. In this method, conformational dynamics is propagated in explicit solvent and lipids, while the solvation forces for propagating titration coordinates are calculated using the membrane generalized-Born GBSW model[S20] based on the conformations sampled in explicit solvent. To accelerate convergence of the coupled conformational and protonation-state sampling, a replica-exchange protocol in the pH space is used.[S18] The membrane-enabled hybrid-solvent CpHMD method[S18,S19] has been validated for p$K_a$ calculations and pH-dependent simulations of transmembrane proteins.[S19,S21,S22] The detailed protocols can be found here[S23]

Three sets of replica-exchange CpHMD simulations were performed, starting from the equilibrated apo active mOR structure, equilibrated fentanyl-mOR complex in the D147-binding mode, and the fentanyl-mOR complex in the H297-binding mode obtained from the WE-HID simulation in which the FEN–H297 distance is less than 3.5 Å. The pH replica-exchange protocol included 16 replicas in the pH range 2.5–9.5 with an increment of 0.5 unit. A GB calculation was invoked every 10 MD steps to update the titration coordinates. In the GB calculation, the default settings were used, consistent with our previous work.[S18] Each pH replica underwent molecular dynamics in the NPT ensemble with an aggregate sampling time of 320 ns. All Asp, Glu, and His sidechains as well as the piperidine amine were allowed to titrate. The model p$K_a$'s of Asp, Glu, and His are 3.8, 4.2, and 6.5, respectively,[S24] while that of the piperidine amine in fentanyl is 8.9.[S25]

**Molecular dynamics protocol.** The temperature and pressure were maintained at 310 K and 1 atm by the Langevin thermostat and Monte Carlo barostat, respectively in the simulations with the Amber program,[S1] while the modified Hoover thermostat and Langevin piston coupling method were used in the simulations with the CHARMM program.[S2] Long-range electrostatics was treated by the particle-mesh Ewald (PME) method[S26] with a real-space cut-off of 12 Å and a sixth-order interpolation with a 1.6-Å$^{-1}$ grid spacing. The van der Waals interactions were smoothly switched to zero between 10 and 12 Å. Bonds involving hydrogen atoms were constrained using the SHAKE algorithm to enable a 2-fs timestep. Analysis was performed using CPPTRAJ program.[S27] For analysis of the equilibrium simulations starting from the D147- and H297-binding modes, the last 200 or 400 ns data was used, respectively.

**Clustering analysis.** The clustering analysis was performed using the cluster command in CPPTRAJ[S27] with the hierarchical agglomerate algorithm. The distance between clusters was calculated based on the RMSD of fentanyl's heavy atoms. The distance cutoff was 3 Å.

**Calculation of Tanimoto coefficients.** Tanimoto coefficient between profiles A and B is calculated using the following equation:

$$S_{AB} = \frac{\sum_{i=1}^{n} x_{iA}.x_{iB}}{\sum_{i=1}^{n}(x_{iA})^2 + \sum_{i=1}^{n}(x_{iB})^2 - \sum_{i=1}^{n} x_{iA}.x_{iB}}$$

where $x_{iA}$ or $x_{iB}$ denotes the contact value between fentanyl and residue $i$ of mOR from profile A or B, respectively. $x_i$ has a value of 1 if a contact exists and 0 otherwise.

**Binding site volume calculation.** A reviewer noted that BU72 is big and so the second binding mode may be the result of using the BU72-bound mOR crystal structure as a template to generate the initial structure for the fentanyl-mOR complex. The reviewer suggested running a 1-$\mu$s MD equilibration of the empty receptor (with the goal to "shrink the binding pocket"). To address this comment, we performed the binding site volume calculations using POVME2.0. [S28,S29] All structures were first aligned using the C$\alpha$ atoms of the binding site residues (Tyr75, Gln124, Asn127, Trp133, Ile144, Asp147, Tyr148, Met151, Phe152, Leu232, Lys233, Val236, Ala240, Trp293, Ile296, His297, Val300, Trp318, His319, Ile322, Tyr326) from the BU72-bound mOR crystal structure (PDB ID: 5C1M). All waters, ions, cholesterol, lipids, nanobody, and ligands were removed before the calculation. The binding pocket searching region is kept consistent throughout all systems by specifying an Inclusion box centered at the binding pocket center of mass (0,0,8) and sides of 12, 12, 15 Å in the x, y, and z direction, respectively. For the volume calculation, trajectory snapshots were taken every 5 ns from the last 50 ns of the simulations.

The binding site volume based on the BU72-bound mOR crystal structure (with BU72 removed) is 371 Å$^3$. After 110 ns of MD equilibration before docking fentanyl, the volume is 359±12 Å$^3$, which is similar to that from the crystal structure. To test if prolonged equilibration would shrink the binding site volume, we extended the simulation by 500 ns and found that the receptor's cavity volume increased to 465±51 Å$^3$. An increase in volume can be rationalized as the result of relaxation and solvation of the binding site. Thus, a long MD equilibration of the empty receptor will not reduce the binding site volume, and the alternative binding mode of fentanyl is not a result of an expanded binding cavity due to the size of BU72.

### 2 Supplementary Tables

Table S1: Steps in the equilibration simulation of apo active mOR

| Step | Ensemble | Time (ns) | BB [a] | mOR Wat[a] | Lipid[b] |
|---|---|---|---|---|---|
| 1 | NVT | 0.35 | 10.0 | 5.00 | 5.0 / 2500 |
| 2 | NVT | 0.15 | 5.00 | 2.50 | 5.0 / 200 |
| 3 | NPT | 0.50 | 5.00 | 2.50 | 5.0 / 200 |
| 4 | NPT | 0.50 | 2.50 | 1.25 | 2.0 / 100 |
| 5 | NPT | 1.00 | 2.50 | 1.25 | 1.0 / 100 |
| 6 | NPT | 1.00 | 2.50 | 1.25 | 0.5 / 50 |
| 7 | NPT | 4.00 | 2.50 | 1.25 | 0 |
| 8 | NPT | 60.0 | 2.50 | 1.25 | 0 |
| 9 | NPT | 50.0 | 0 | 0 | 0 |

[a] Force constant in the harmonic restraint potential for the backbone heavy atoms in kcal/mol/Å$^2$. [b] Force constant in the position/dihedral restraint potential. The unit is kcal/mol/Å$^2$ for the position restraints or kcal/mol for dihedral angle restraints. mOR Wat referes to the crystal waters and seven additional water added using the program DOWSER.[S10]

Table S2: Steps in the equilibration simulation to relax the docked fentanyl-mOR complex structure

| Step | Ensemble | Time (ns) | BB[a] | SC[a] | fentanyl[b] |
|---|---|---|---|---|---|
| 1 | NPT | 1.00 | 2.50 | 1.25 | 2.50 |
| 2 | NPT | 1.00 | 1.25 | 0.75 | 1.25 |
| 3 | NPT | 1.00 | 1.00 | 0.50 | 1.00 |
| 4 | NPT | 1.00 | 0.50 | 0.25 | 0.50 |
| 5 | NPT | 1.00 | 0.25 | 0 | 0.25 |
| 6 | NPT | 110.0 | 0 | 0 | 0 |

[a] Force constant in the harmonic restraint potential for the backbone and sidechain heavy atoms in kcal/mol/Å$^2$. [b] Force constant in the harmonic restraint potential for the fentanyl heavy atoms.

Table S3: Steps in the equilibration simulation to relax the H297 binding mode with H297 as either HIE or HIP

| Step | Ensemble | Time (ns) | BB[a] | SC[a] | fentanyl[b] |
|---|---|---|---|---|---|
| 1 | NPT | 1.00 | 2.50 | 1.25 | 2.50 |
| 2 | NPT | 1.00 | 1.25 | 0.75 | 1.25 |
| 3 | NPT | 1.00 | 1.00 | 0.50 | 1.00 |
| 4 | NPT | 1.00 | 0.50 | 0.25 | 0.50 |
| 5 | NPT | 1.00 | 0.25 | 0 | 0.25 |
| 6 | NPT | 60.0 | 0 | 0 | 0 |

[a] Force constant in the harmonic restraint potential for the backbone and sidechain heavy atoms in kcal/mol/Å$^2$. [b] Force constant in the harmonic restraint potential for the fentanyl heavy atoms.

Table S4: Calculated p$K_a$'s of mOR titratable residues and fentanyl from the replica-exchange CpHMD simulations

| Residue | CpH-apo | CpH-D147 | CpH-H297 |
|---|---|---|---|
| Asp114 | 4.8 | 5.1 | 4.8 |
| Asp147 | 3.3 | 4.1 | 3.8 |
| Asp164 | 3.6 | 2.8 | 3.0 |
| Asp177 | 3.6 | 3.8 | 3.8 |
| Asp216 | 2.5 | 2.9 | 2.5 |
| Asp272 | 2.6 | 2.0 | 1.9 |
| Asp340 | 3.3 | 3.3 | 3.2 |
| Glu229 | 3.2 | 2.8 | 2.9 |
| Glu270 | 3.5 | 3.3 | 3.3 |
| Glu310 | 3.8 | 3.2 | 3.7 |
| Glu341 | 3.2 | 3.5 | 3.0 |
| His54 | 6.8 | 7.3 | 7.5 |
| His171 | 5.6 | 5.3 | 5.2 |
| His223 | 5.8 | 5.8 | 5.7 |
| His297 | 6.8 | 7.3 | 6.7 |
| His319 | 7.1 | 7.2 | 6.8 |
| Fentanyl | NA | >9.5[a] | >9.5[a] |

[a] Only a lower bound to the p$K_a$ of the piperidine amine of fentanyl is given, as the amine remains charged in the entire simulation pH range 2.5–9.5.

### 3 Supplementary Figures

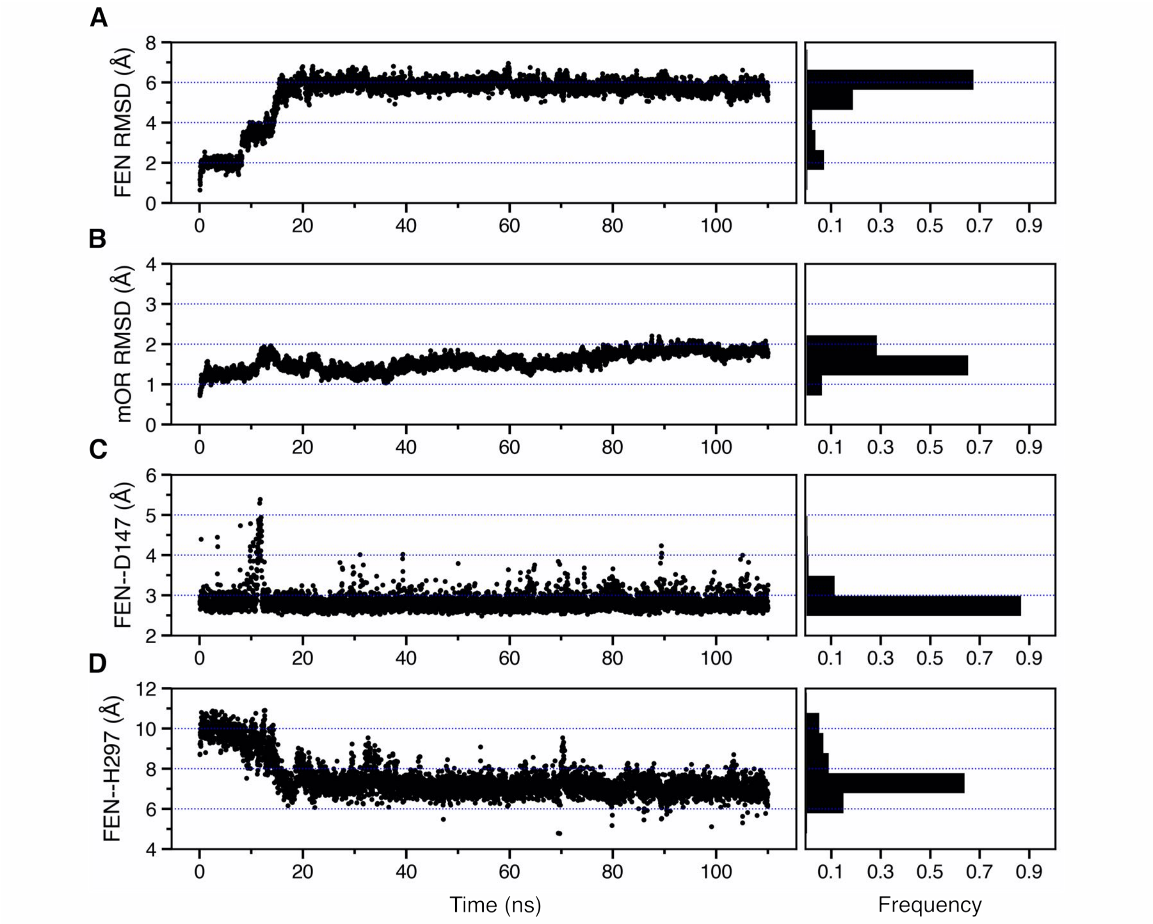


Figure S1: **Equilibration of the docked fentanyl-mOR structure. A, B.** Time series of the root-mean-square deviation (RMSD) of the fentanyl **(A)** and mOR **(B)** heavy atom positions with respect to their starting positions. **C.** Time series of the FEN–D147 distance, defined as the minimum distance between the piperidine nitrogen and the carboxylate oxygen of Asp147. **D.** Time series of the FEN–H297 distance, defined as that between the piperidine nitrogen and the unprotonated imidazole nitrogen of His297.

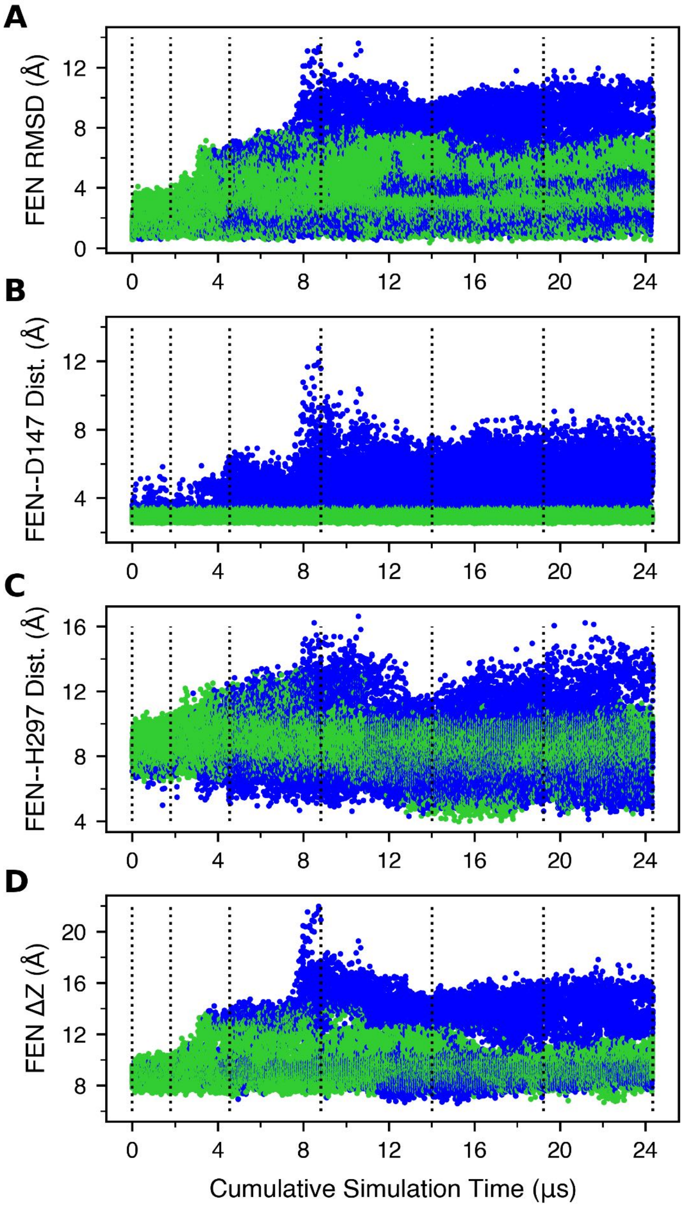


Figure S2: **Characterization of the fentanyl position as a function of the cumulative simulation time from the WE-HIE simulation. A.** The root-mean-square deviation (RMSD) of the fentanyl heavy atoms with respect to the starting position. **B.** The FEN–D147 distance refers to the minimum distance between the piperidine nitrogen and the carboxylate oxygen of Asp147. **C.** The FEN–H297 distance is measured between the piperidine nitrogen and the unprotonated imidazole nitrogen of His297. **D.** Fentanyl $\Delta Z$ position is defined as the distance between the centers of mass (COM) of fentanyl and mOR in the z direction. Data with the FEN–D147 and FEN–H297 distances below 3.5 Å are colored green and orange, respectively, and otherwise blue. The unweighted data from all bins were taken and the time refers to the cumulative time. the vertical lines are drawn at every 50 weighted ensemble iterations.

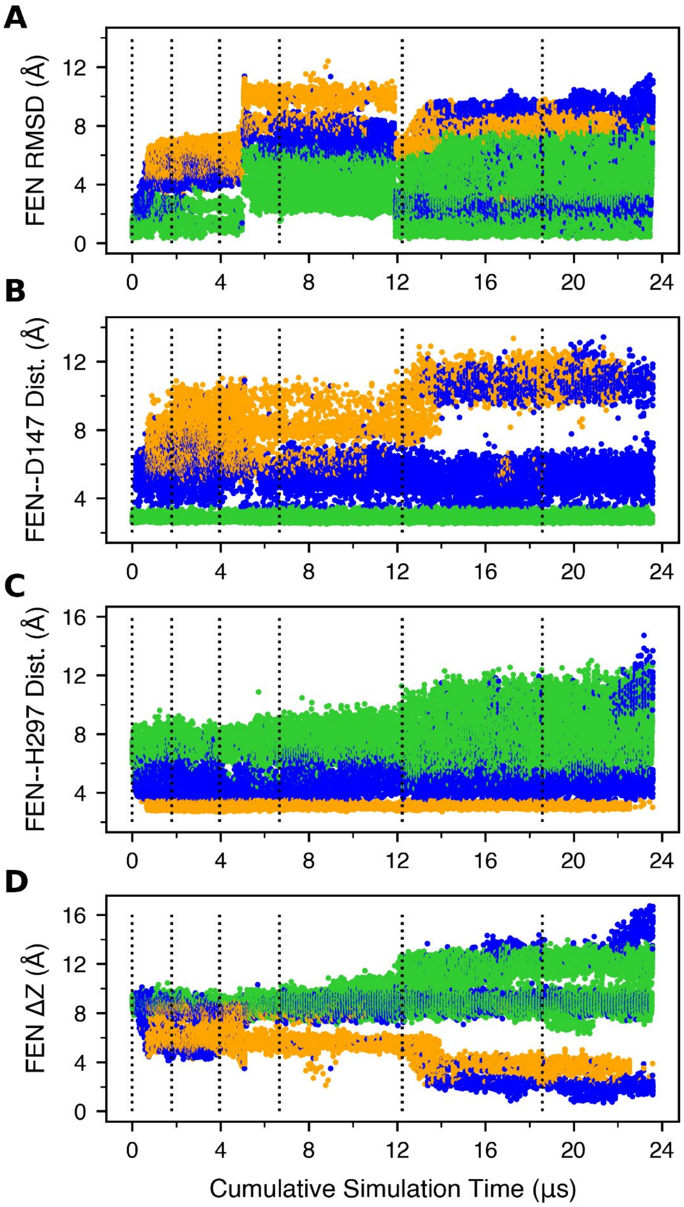


Figure S3: **Characterization of the fentanyl position as a function of the cumulative simulation time from the WE-HID simulation. A.** The root-mean-square deviation (RMSD) of the fentanyl heavy atoms. An abrupt change near $\sim$5 $\mu$s is due to the manual bin boundary modification and does not affect the sampling results. **B.** The FEN–D147 distance refers to the minimum distance between the piperidine nitrogen and the carboxylate oxygen of Asp147. **C.** The FEN–H297 distance is measured between the piperidine nitrogen and the unprotonated imidazole nitrogen of His297. **D.** Fentanyl $\Delta$Z position is defined as the distance between the centers of mass (COM) of fentanyl and mOR in the z direction. Data with the FEN–D147 and FEN–H297 distances below 3.5 Å are colored green and orange, respectively, and otherwise blue. The unweighted data from all bins were taken and the time refers to the cumulative time. the vertical lines are drawn at every 50 weighted ensemble iterations.

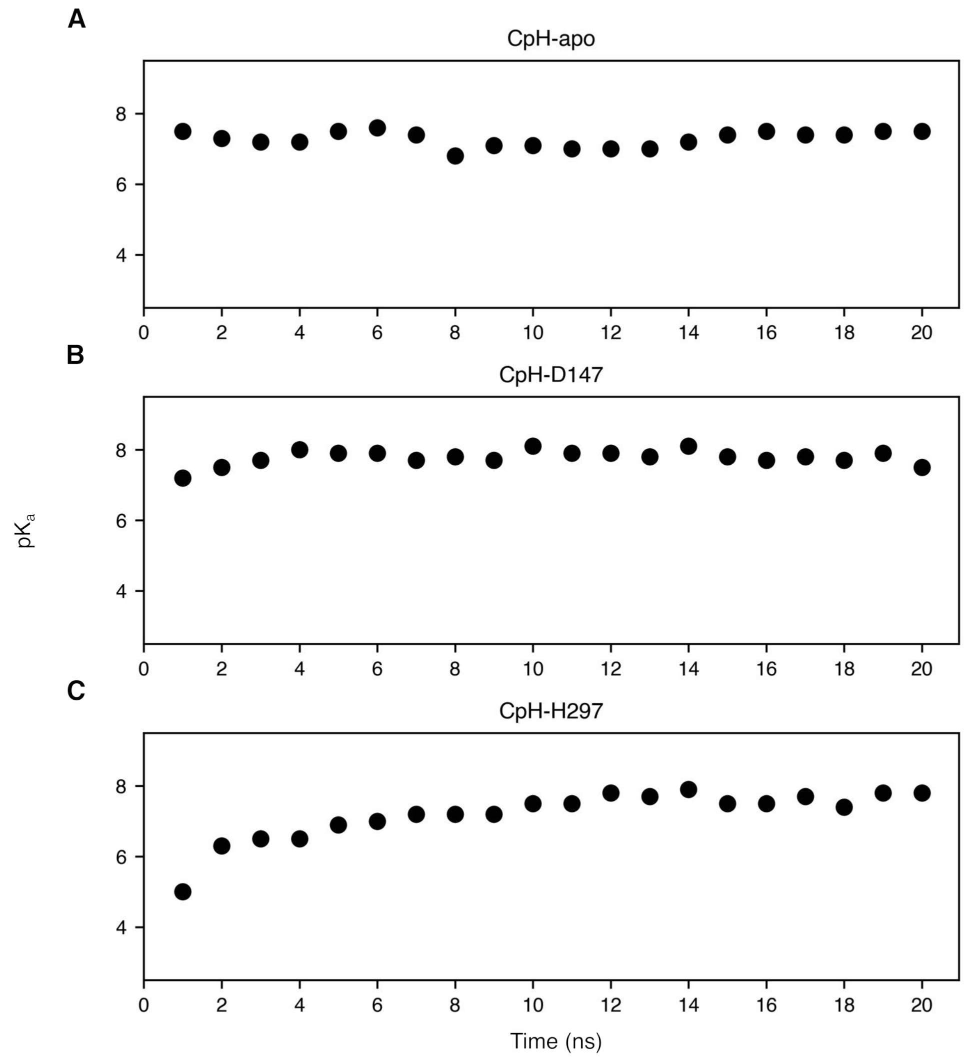


Figure S4: **The p$K_a$ of His297 is well converged in all three sets of replica-exchange CpHMD simulations.** The p$K_a$ value of His297 plotted every 1-ns as a function of the single replica sampling time from the CpH-apo **(A)**, CpH-D147 **(B)**, and CpH-H297 **(C)** simulations.

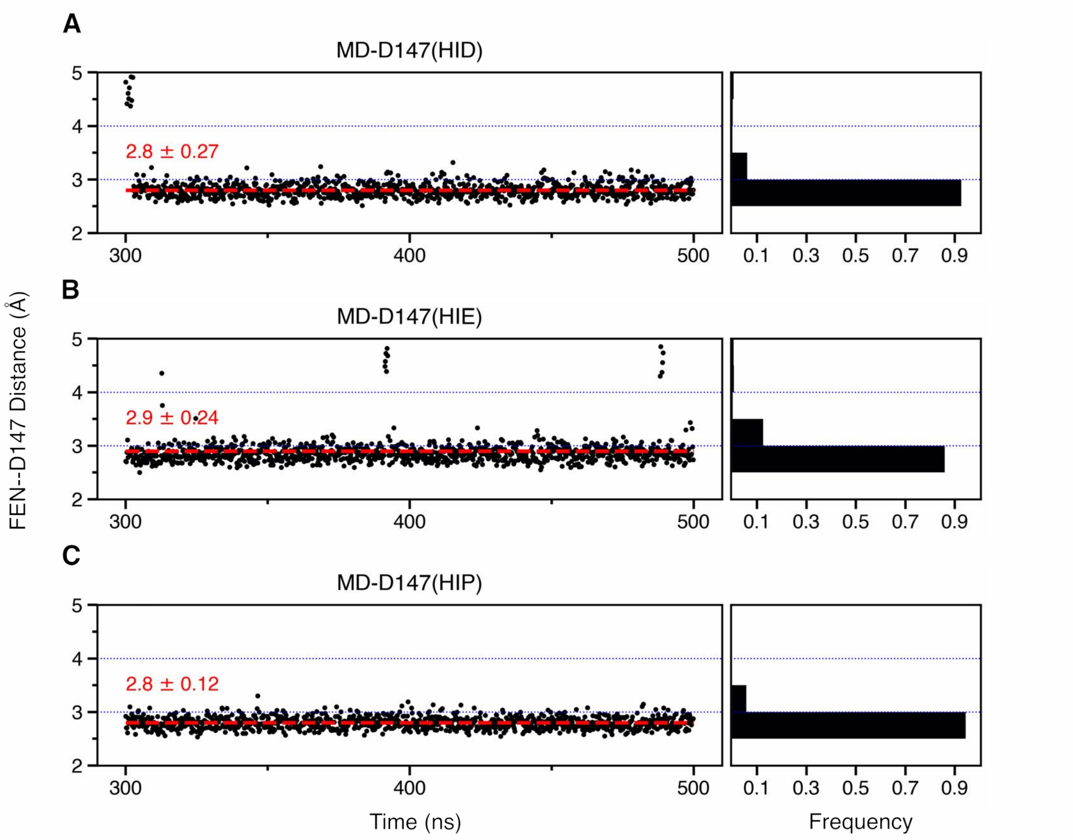


Figure S5: **The fentanyl-D147 salt bridge remains stable with all three protonation states of His297 in the equilibrium simulations of the D147-binding mode.** The time series of the FEN–D147 distance in the presence of HID297 **(A)**, HIE297 **(B)** , and HIP297 **(C)**. The FEN–D147 distance refers to the minimum distance between the piperidine nitrogen and the carboxylate oxygen of Asp147.

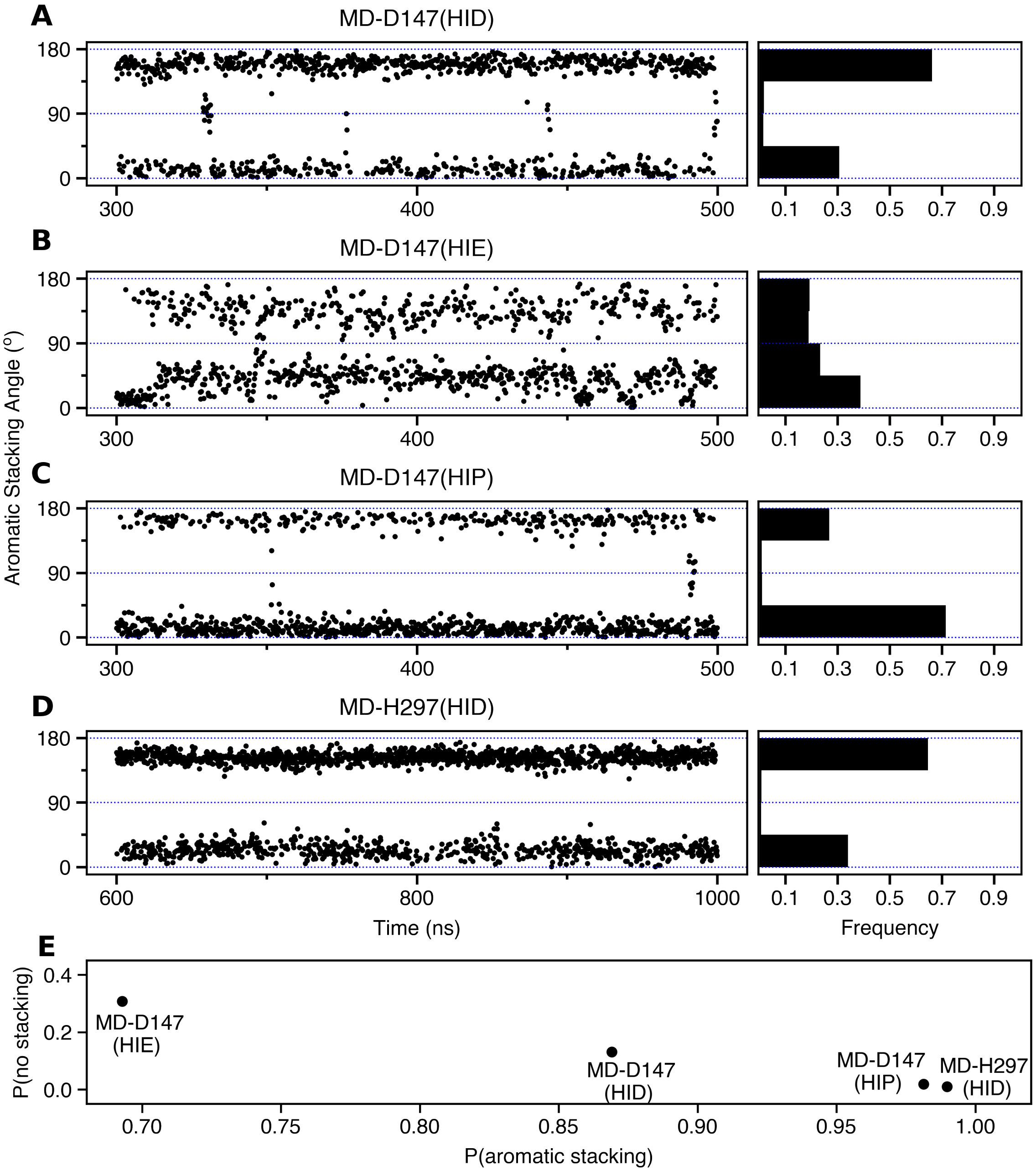


Figure S6: **Aromatic stacking interaction between the phenyl ring of fentanyl's phenethyl group and Trp293 is persistent for both D147- and H297-binding modes**. Time series of the fentanyl-W293 stacking angle from the D147-binding-mode simulations with HID297 **(A)**, HIE297 **(B)**, or HIP297 **(C)**, and from the H297-binding-mode simulation with HID297 **(D)**. **E.** Relationship between the probabilities of aromatic stacking and no aromatic stacking from each simulation. The angle is measured between the planes of the phenyl ring of fentanyl's phenethyl group and the six-member ring of Trp293. There is aromatic stacking interaction if the angle between the two planes is within 0–45° or 135–180°.

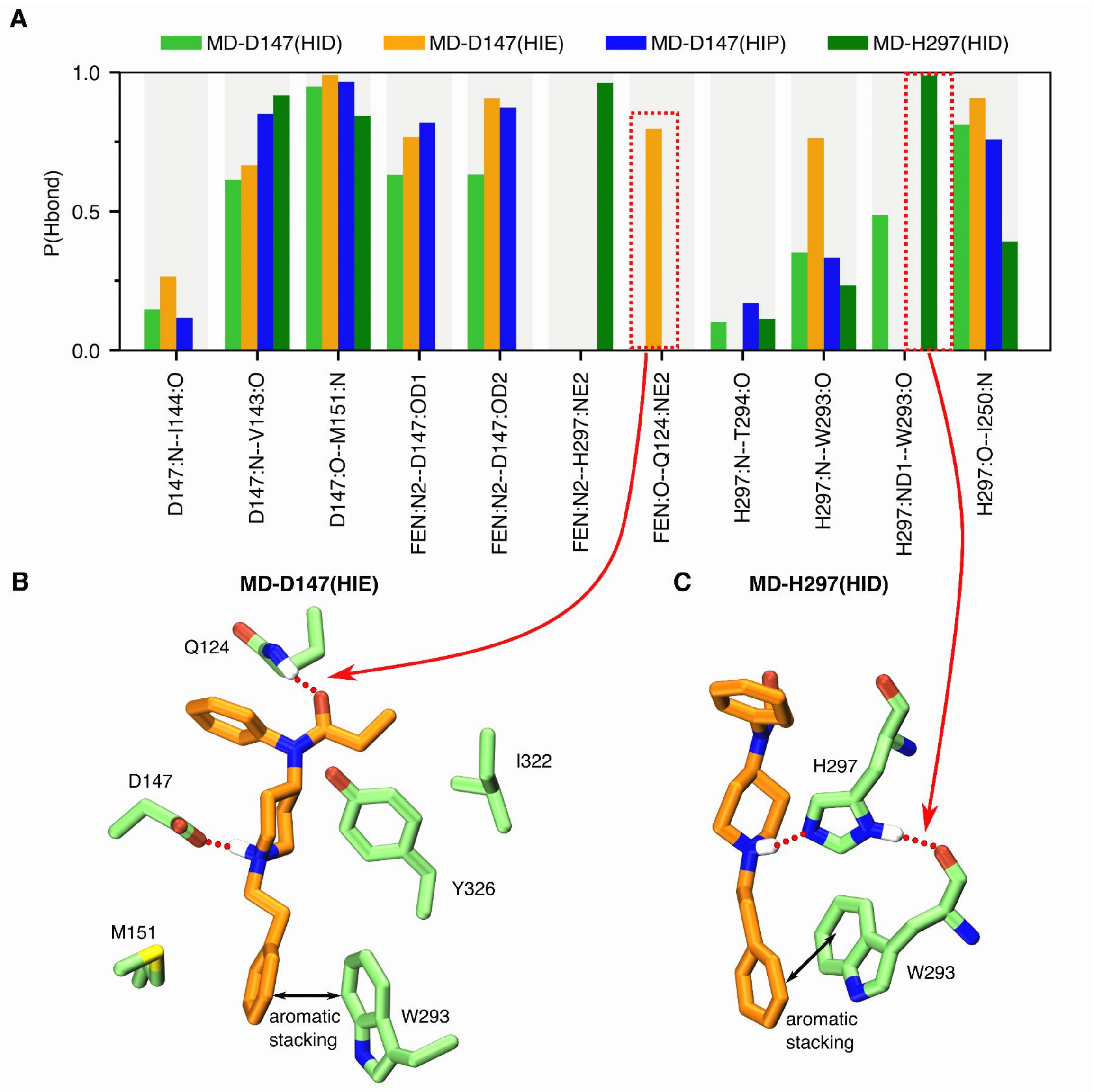


Figure S7: **Occupancies of the hydrogen bonds formed between fentanyl-mOR and mOR-mOR from the simulations of the D147- and H297-binding modes.** The bars are colored as shown in top legend. A value of 1 means the hydrogen bond is persistent for the entire duration of the trajectory. A hydrogen bond is considered present if the donor-acceptor distance is below 3.5 Å and the acceptor-H-donor angle is less than 120°.

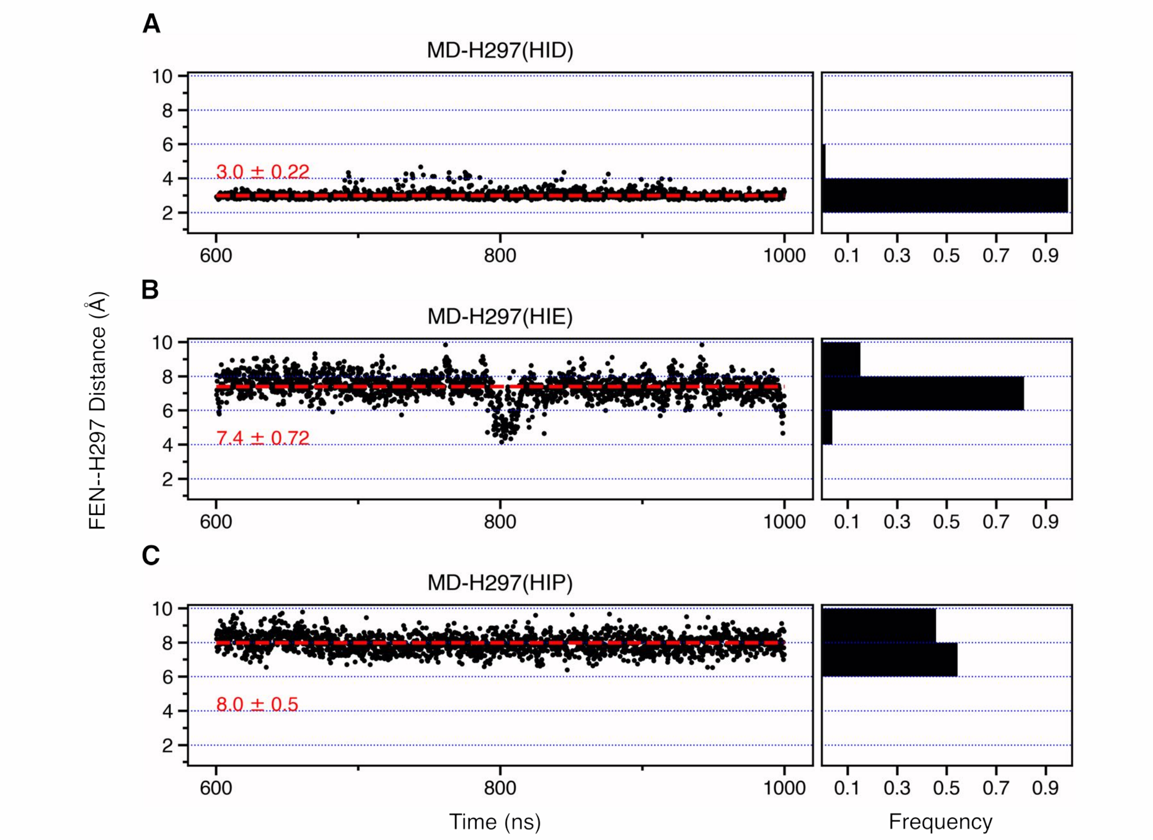


Figure S8: **Fentanyl is pushed away from His297 in the equilibrium simulations of the H297-binding mode with HIE and HIP.** The FEN–H297 distance as a function of simulation time in H297 mode with HID297 **(A)**, HIE297 **(B)**, and HIP297 **(C)**. The FEN–H297 distance is measured between the piperidine nitrogen and the unprotonated imidazole nitrogen of His297.

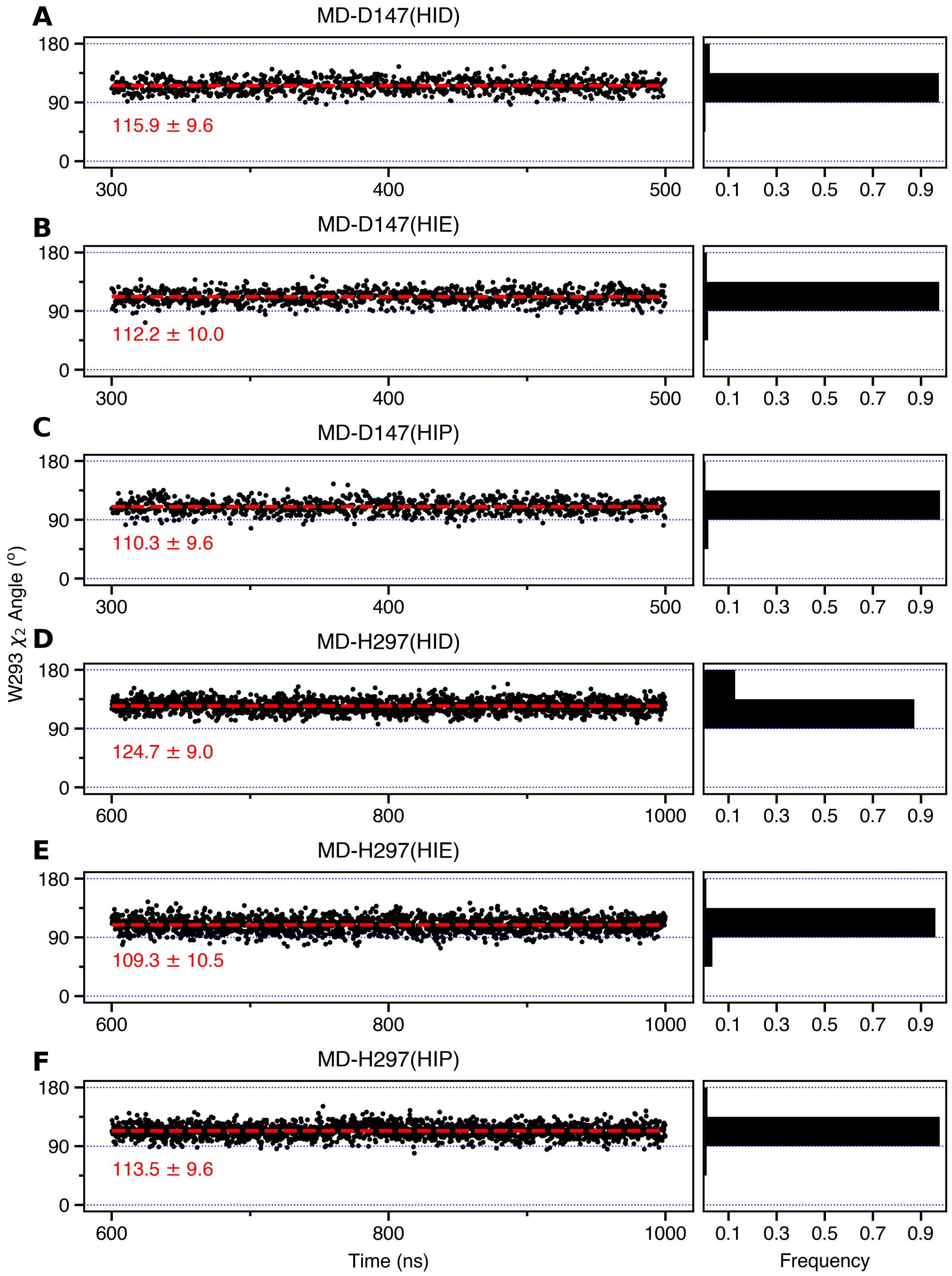


Figure S9: **Trp293 $\chi_2$ angle is somewhat decrease or increased relative to the value (120°) in the crystal structure of BU72-bound mOR (PDB 5C1M).[S9] Time series of the Trp293 $\chi_2$ angle in the simulations of the D147- (A-C) and the H297-binding mode (D-F).**

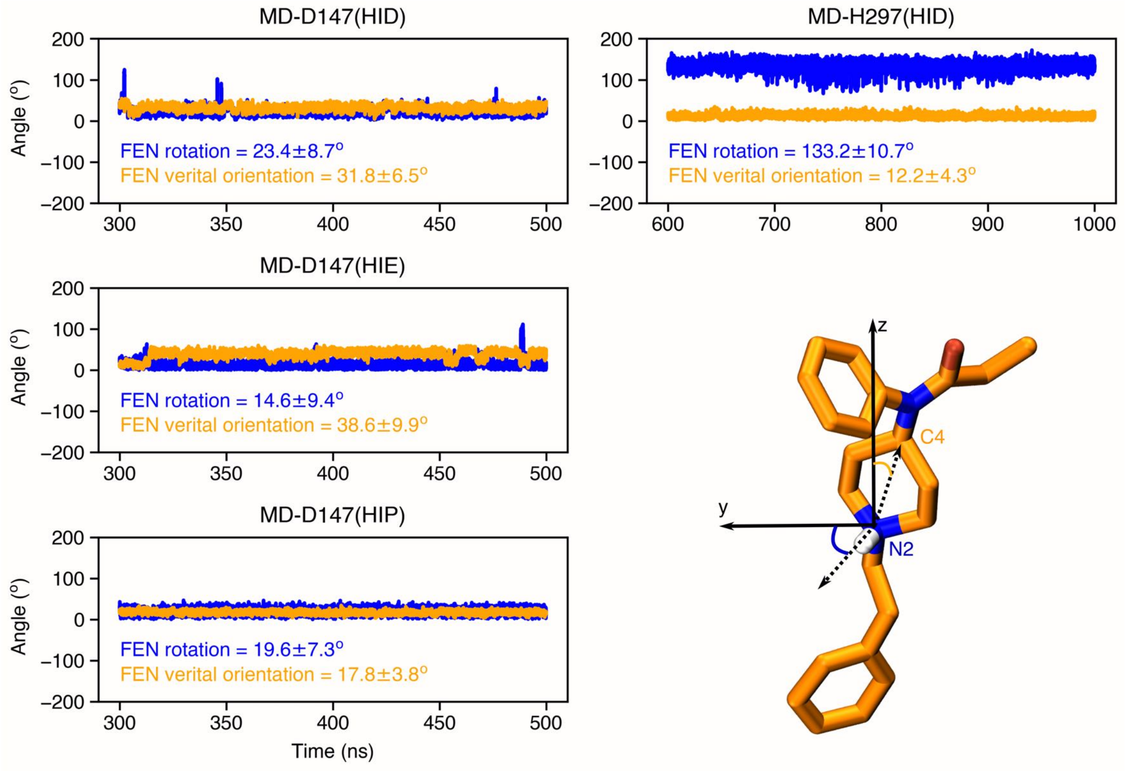


Figure S10: **Fentanyl adopts different orientations in the simulations of the D147- and H297-binding modes**. Time series of the fentanyl rotation angle (blue) and vertical orientation angle (gold). Fentanyl rotation angle is measured by the angle between N2–H vector and the y-axis. Fentanyl vertical orientation angle is measured by the angle between the N2–C4 vector and the membrane normal (z-axis).

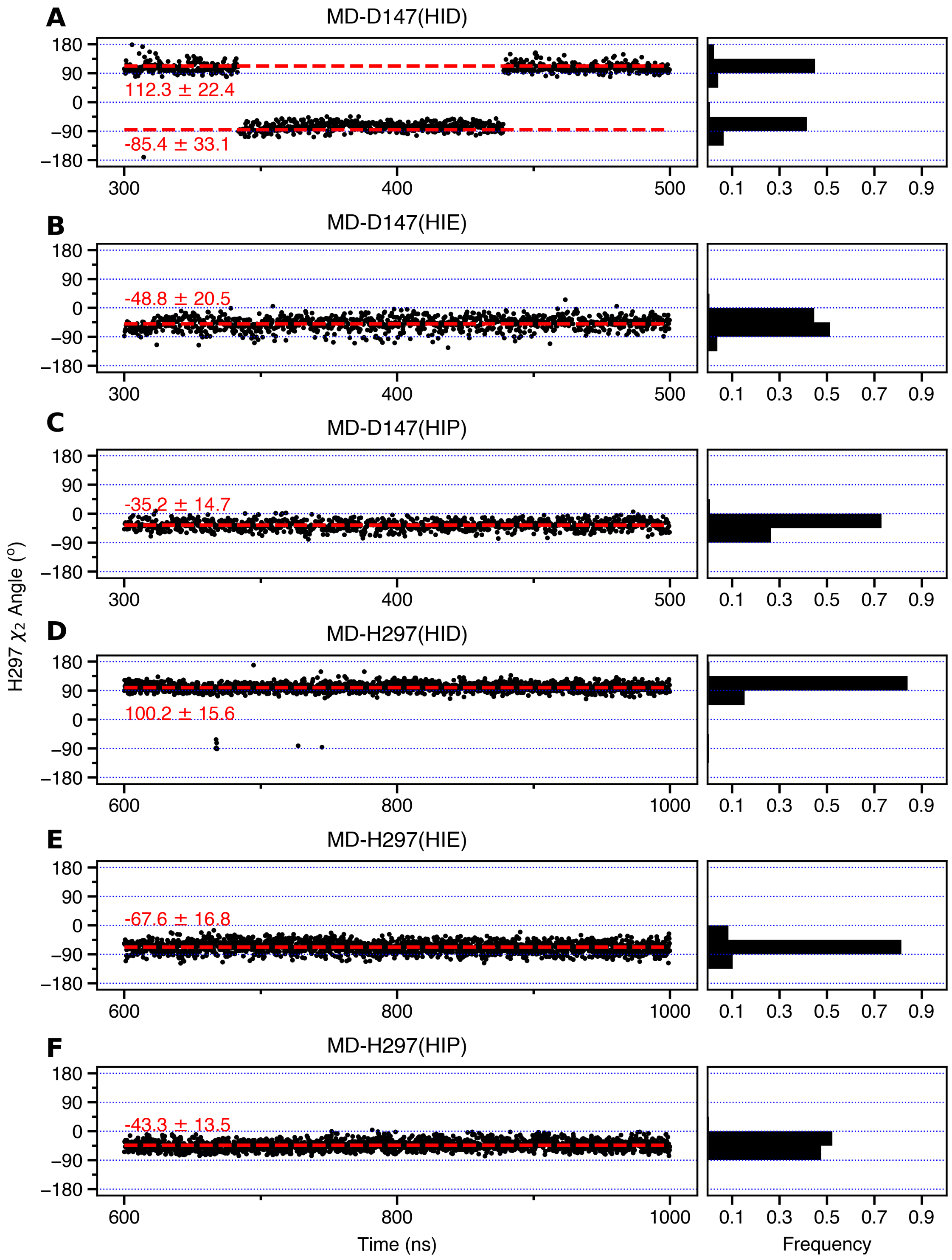


Figure S11: $\chi_2$ **angle of His297 appears to be influenced by the protonation/tautomer state.** Time series of the $\chi_2$ angle of His297 in the simulations of the D147-binding mode (**A-C**) and the H297-binding mode (**D-F**).

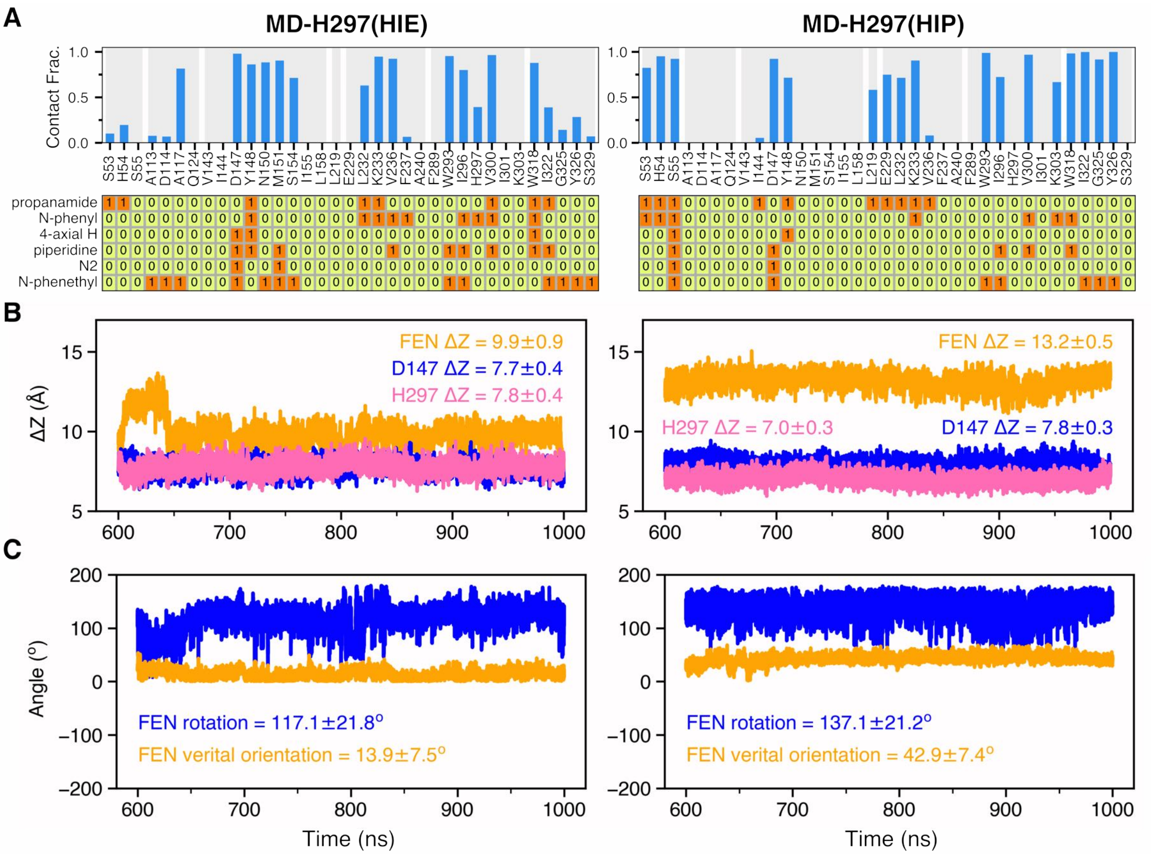


Figure S12: **Characterization of the equilibrium simulations of the H297-binding mode with HIE297 and HIP297.** MD-H297(HIE) (left column) and MD-H297(HIP) (right column) simulations. **A.** Fentanyl-mOR contact profile (top) and fingerprint matrix (bottom) from the simulations with HIE297 (left) and HIP297 (right). Details are explained in the caption of Fig. 4 of the main text. **B.** Time series of the vertical positions of fentanyl (gold), Asp147 (blue), and His297 (magenta) from the simulations with HIE297 (left) and HIP297 (right). **C.** Time series of the fentanyl rotation angle (blue) and vertical orientation angle (gold) from the simulations with HIE297 (left) and HIP297 (right). Fentanyl rotation angle is measured by the angle between N2–H vector and the y-axis. Fentanyl vertical orientation angle is measured by the angle between the N2–C4 vector and the membrane normal (z-axis).

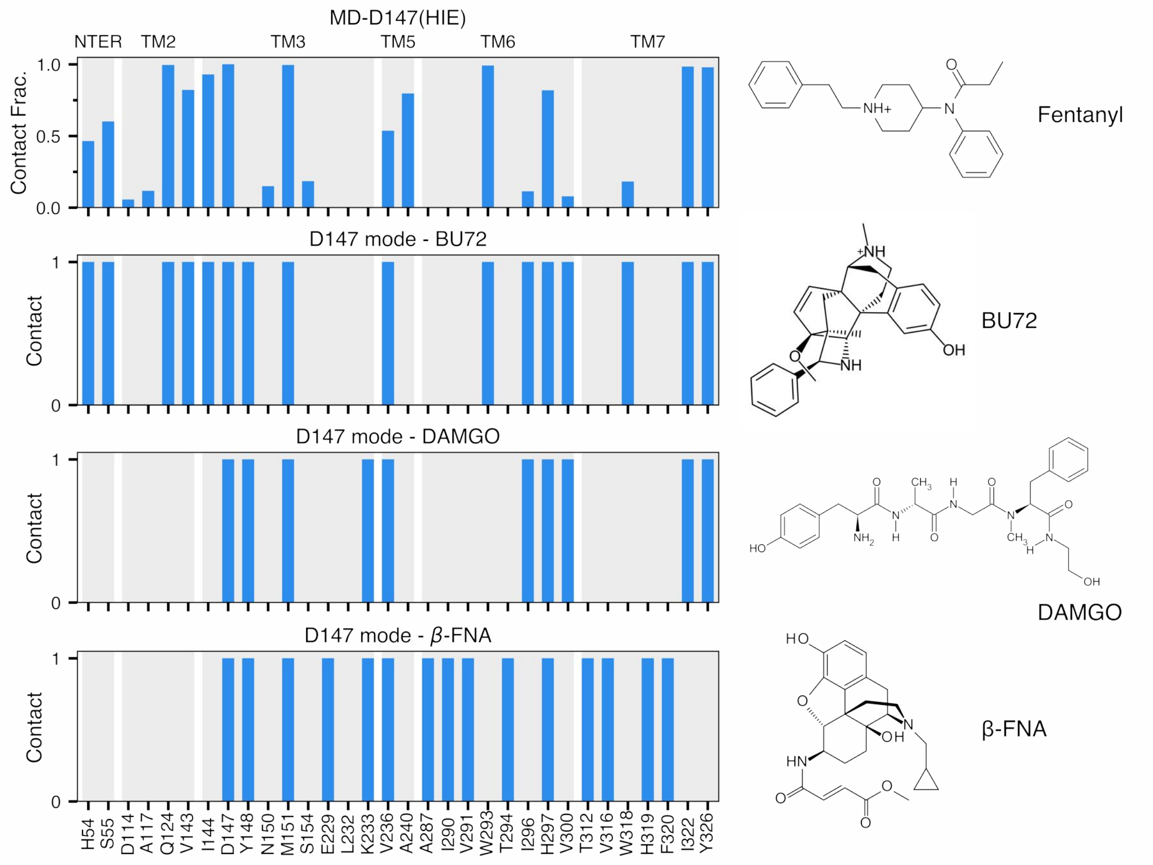


Figure S13: **Comparison between fentanyl-mOR contact fraction to the BU72-, DAMGO-, $\beta$-FNA-mOR contacts in the crystal structures.** MD-D147(HIE) was shown due to its highest $T_c$ value compared to BU72 (Figure 5 in main). The N-terminal residues (residue 52 to 64) are not resolved in the crystal structures of DAMGO- and $\beta$-FNA-mOR.

#### References


(S1) Case, D. A.; Ben-Shalom, I. Y.; Brozell, S. R.; Cerutti, D. S.; Cheatham, T. E. I.; Cruzeiro, V. W. D.; Darden, T. A.; Duke, R. E.; Ghoreishi, D.; Gilson, M. K. AMBER 2018. 2018.

(S2) Brooks, B. R.; Brooks, C. L.; Mackerell, A. D.; Nilsson, L.; Petrella, R. J.; Roux, B.; Won, Y.; Archontis, G.; Bartels, C.; Boresch, S.; Caflisch, A.; Caves, L.; Cui, Q.; Dinner, A. R.; Feig, M.; Fischer, S.; Gao, J.; Hodoscek, M.; Im, W.; Kuczera, K.; Lazaridis, T.; Ma, J.; Ovchinnikov, V.; Paci, E.; Pastor, R. W.; Post, C. B.; Pu, J. Z.; Schaefer, M.; Tidor, B.; Venable, R. M.; Woodcock, H. L.; Wu, X.; Yang, W.; York, D. M.; Karplus, M. CHARMM: The Biomolecular Simulation Program. *J. Comput. Chem.* **2009**, *30*, 1545–1614.

(S3) Best, R. B.; Zhu, X.; Shim, J.; Lopes, P. E. M.; Mittal, J.; Feig, M.; MacKerell, A. D. Optimization of the Additive CHARMM All-Atom Protein Force Field Targeting Improved Sampling of the Backbone $\phi$, $\psi$ and Side-Chain $\chi_1$ and $\chi_2$ Dihedral Angles. *J. Chem. Theory Comput.* **2012**, *8*, 3257–3273.

(S4) MacKerell, Jr., A. D.; Bashford, D.; Bellott, M.; Dunbrack, Jr., R. L.; Evanseck, J. D.; Field, M. J.; Fischer, S.; Gao, J.; Guo, H.; Ha, S.; Joseph-McCarthy, D.; Kuchnir, L.; Kuczera, K.; Lau, F. T. K.; Mattos, C.; Michnick, S.; Ngo, T.; Nguyen, D. T.; Prodhom, B.; Reiher, III, W. E.; Roux, B.; Schlenkrich, M.; Smith, J. C.; Stote, R.; Straub, J.; Watanabe, M.; Wiórkiewicz-Kuczera, J.; Yin, D.; Karplus, M. All-atom empirical potential for molecular modeling and dynamics studies of proteins. *J. Phys. Chem. B* **1998**, *102*, 3586–3616.

(S5) MacKerell, Jr., A. D.; Feig, M.; Brooks, III, C. L. Extending the treatment of backbone energetics in protein force fields: limitations of gas-phase quantum mechanics in reproducing protein conformational distributions in molecular dynamics simulations. *J. Comput. Chem.* **2004**, *25*, 1400–1415.

(S6) Klauda, J. B.; Venable, R. M.; Freites, J. A.; O'Connor, J. W.; Tobias, D. J.; Mondragon-Ramirez, C.; Vorobyov, I.; MacKerell, A. D.; Pastor, R. W. Update of the CHARMM All-Atom Additive Force Field for Lipids: Validation on Six Lipid Types. *J. Phys. Chem. B* **2010**, *114*, 7830–7843.

(S7) Lim, J. B.; Rogaski, B.; Klauda, J. B. Update of the Cholesterol Force Field Parameters in CHARMM. *J. Phys. Chem. B* **2012**, *116*, 203–210.

(S8) Vanommeslaeghe, K.; Raman, E. P.; MacKerell, A. D. Automation of the CHARMM General Force Field (CGenFF) II: Assignment of Bonded Parameters and Partial Atomic Charges. *J. Chem. Inf. Model.* **2012**, *52*, 3155–3168.

(S9) Huang, W.; Manglik, A.; Venkatakrishnan, A. J.; Laeremans, T.; Feinberg, E. N.; Sanborn, A. L.; Kato, H. E.; Livingston, K. E.; Thorsen, T. S.; Kling, R. C.; Granier, S.; Gmeiner, P.; Husbands, S. M.; Traynor, J. R.; Weis, W. I.; Steyaert, J.; Dror, R. O.; Kobilka, B. K. Structural Insights into $\mu$-Opioid Receptor Activation. *Nature* **2015**, *524*, 315–321.

(S10) Zhang, L.; Hermans, J. Hydrophilicity of Cavities in Proteins. *Proteins* **1996**, *24*, 433.

(S11) Lomize, M. A.; Pogozheva, I. D.; Joo, H.; Mosberg, H. I.; Lomize, A. L. OPM Database and PPM Web Server: Resources for Positioning of Proteins in Membranes. *Nucleic Acids Res.* **2012**, *40*, D370–D376.

(S12) Lee, J.; Cheng, X.; Swails, J. M.; Yeom, M. S.; Eastman, P. K.; Lemkul, J. A.; Wei, S.; Buckner, J.; Jeong, J. C.; Qi, Y.; Jo, S.; Pande, V. S.; Case, D. A.; Brooks, C. L.; MacKerell, A. D.; Klauda, J. B.; Im, W. CHARMM-GUI Input Generator for NAMD, GROMACS, AMBER, OpenMM, and CHARMM/OpenMM Simulations Using the CHARMM36 Additive Force Field. *J. Chem. Theory Comput.* **2016**, *12*, 405–413.

(S13) Ellis, C. R.; Kruhlak, N. L.; Kim, M. T.; Hawkins, E. G.; Stavitskaya, L. Predicting Opioid Receptor Binding Affinity of Pharmacologically Unclassified Designer Substances Using Molecular Docking. *PLoS One* **2018**, *13*, e0197734.

(S14) Lipiński, P. F. J.; Jarończyk, M.; Dobrowolski, J. C.; Sadlej, J. Molecular Dynamics of Fentanyl Bound to $\mu$-Opioid Receptor. *J. Mol. Model.* **2019**, *25*, 144.

(S15) Huber, G.; Kim, S. Weighted-Ensemble Brownian Dynamics Simulations for Protein Association Reactions. *Biophys. J.* **1996**, *70*, 97–110.

(S16) Zwier, M. C.; Kaus, J. W.; Chong, L. T. Efficient Explicit-Solvent Molecular Dynamics Simulations of Molecular Association Kinetics: Methane/Methane, Na+/Cl-, Methane/Benzene, and K+/18-Crown-6 Ether. *J. Chem. Theory Comput.* **2011**, *7*, 1189–1197.

(S17) Zuckerman, D. M.; Chong, L. T. Weighted Ensemble Simulation: Review of Methodology, Applications, and Software. *Annu. Rev. Biophys.* **2017**, *46*, 43–57.

(S18) Wallace, J. A.; Shen, J. K. Continuous Constant pH Molecular Dynamics in Explicit Solvent with pH-Based Replica Exchange. *J. Chem. Theory Comput.* **2011**, *7*, 2617–2629.

(S19) Huang, Y.; Chen, W.; Dotson, D. L.; Beckstein, O.; Shen, J. Mechanism of pH-Dependent Activation of the Sodium-Proton Antiporter NhaA. *Nat. Commun.* **2016**, *7*, 12940.

(S20) Im, W.; Feig, M.; Brooks, C. L. An Implicit Membrane Generalized Born Theory for the Study of Structure, Stability, and Interactions of Membrane Proteins. *Biophys. J.* **2003**, *85*, 2900–2918.

(S21) Chen, W.; Huang, Y.; Shen, J. Conformational Activation of a Transmembrane Proton Channel from Constant pH Molecular Dynamics. *J. Phys. Chem. Lett.* **2016**, *7*, 3961–3966.

(S22) Yue, Z.; Chen, W.; Zgurskaya, H. I.; Shen, J. Constant pH Molecular Dynamics Reveals How Proton Release Drives the Conformational Transition of a Transmembrane Efflux Pump. *J. Chem. Theory Comput.* **2017**, *13*, 6405–6414.

(S23) Huang, Y.; Henderson, J. A.; Shen, J. Continuous constant pH Molecular Dynamics Simulations of Transmembrane Proteins. *Methods Mol. Biol.* **2020**, in press.

(S24) Thurlkill, R. L.; Grimsley, G. R.; Scholtz, J. M.; Pace, C. N. pK values of the ionizable groups of proteins. *Protein Sci.* **2006**, *15*, 1214–1218.

(S25) Roy, S. D.; Flynn, G. L. Solubility Behavior of Narcotic Analgesics in Aqueous Media: Solubilities and Dissociation Constants of Morphine, Fentanyl, and Sufentanil. *Pharm. Res.* **1989**, *6*, 147–151.

(S26) Darden, T.; York, D.; Pedersen, L. Particle Mesh Ewald: An $N\cdot\log(N)$ Method for Ewald Sums in Large Systems. *J. Chem. Phys.* **1993**, *98*, 10089–10092.

(S27) Roe, D. R.; Cheatham, T. E. PTRAJ and CPPTRAJ: Software for Processing and Analysis of Molecular Dynamics Trajectory Data. *J. Chem. Theory Comput.* **2013**, *9*, 3084–3095.

(S28) Durrant, J. D.; de Oliveira, C. A. F.; McCammon, J. A. POVME: An Algorithm for Measuring Binding-Pocket Volumes. *J. Mol. Graph. Model.* **2011**, *29*, 773–776.

(S29) Durrant, J. D.; Votapka, L.; Sørensen, J.; Amaro, R. E. POVME 2.0: An Enhanced

Tool for Determining Pocket Shape and Volume Characteristics. *J. Chem. Theory Comput.* **2014**, *10*, 5047–5056.
